# Analysis of copy number variants on chromosome 21 in Down syndrome-associated congenital heart defects

**DOI:** 10.1101/093583

**Authors:** Benjamin L. Rambo-Martin, Jennifer G. Mulle, David J. Cutler, Lora J.H. Bean, Tracie C. Rosser, Kenneth J. Dooley, Clifford Cua, George Capone, Cheryl L. Maslen, Roger H. Reeves, Stephanie L. Sherman, Michael E. Zwick

## Abstract

One in five people with Down syndrome (DS) are born with an atrioventricular septal defect (AVSD), an incidence 2,000 times higher than in the euploid population. The genetic loci that contribute to this risk are poorly understood. In this study, we tested two hypotheses: 1) individuals with DS carrying chromosome 21 copy number variants (CNVs) that interrupt exons may be protected from AVSD, because these CNVs return AVSD susceptibility loci back to disomy, and 2) individuals with DS carrying chromosome 21 genes spanned by microduplications are at greater risk for AVSD because these microduplications boost the dosage of AVSD susceptibility loci beyond a tolerable threshold. We tested 198 case individuals with DS+AVSD and 211 control individuals with DS and a normal heart using a custom microarray with dense probes tiled on chromosome 21 for array CGH. We found that neither an individual chromosome 21 CNV nor any individual gene intersected by a CNV was associated with AVSD in DS. Burden analyses revealed that African American controls had more bases covered by rare deletions than did African American cases. Inversely, we found that Caucasian cases had more genes intersected by rare duplications than did Caucasian controls. Pathway analyses indicated copy number perturbations of genes involved in protein heterotrimerization and histone methylating proteins. Finally, we showed that previously DS+AVSD-associated common CNVs on chromosome 21 are likely false positives. This research adds to the swell of evidence indicating that DS-associated AVSD is similarly heterogeneous, as is AVSD in the euploid population.

## Introduction

Understanding the rules by which variation that influences genome dosage also impacts phenotypes remains one of the central challenges of human genetics (Zarrei et al. 2015). Down syndrome (DS), caused largely by trisomy 21, provides an extreme example of a dosage change that impacts many aspects of an individual’s phenotype. Congenital heart defects (CHDs) are among the most common and significant birth defects found in individuals with DS. In the disomic population, CHDs are the most common birth defect, presenting in 80 out of 1,000 live births and causing 25% of infant mortality (Reller et al. 2008; Yang et al. 2006; Hartman et al. 2011; Mai et al. 2015). For children with trisomy 21, CHD incidence is substantially higher: nearly 450 out of 1,000 live births have a CHD (Loffredo et al. 2001; Freeman et al. 2008).

Atrioventricular septal defects (AVSDs) are a serious CHD resulting from the failure of endocardial cushion formation and subsequent mitral and tricuspid valve formation. In the presence of an AVSD, there is improper mixing of oxygenated and deoxygenated blood. While the heart is typically repaired during the first year of life, patients with AVSD face increased risk of sequelae, including arrhythmias, endocarditis, stroke, congestive heart failure, pulmonary hypertension, and continued heart valve problems (Le Gloan et al. 2011). Trisomy 21 is the single greatest risk factor for AVSD. While 1 in 10,000 people in the general population present with AVSD, among infants with DS, the rate is 1 in 5 (Freeman et al. 2008). This 2,000-fold greater risk suggests that those with DS may represent a sensitized population in which genetic variation contributing to the risk for AVSD may have a larger effect size than in the general population. Using the DS population to identify AVSD risk loci may therefore yield high statistical power, even with a small sample size (Zwick et al. 1999).

In our prior study, the largest genetic study of its kind to date, we characterized genome-wide copy number variants (CNVs) in a well-phenotyped cohort with 210 case individuals with DS and a complete AVSD (DS+AVSD) and 242 control individuals with DS and structurally normal hearts (DS+NH) (Ramachandran et al. 2015). We showed a statistically significant increase in large, rare deletions in DS+AVSD cases that also impacted more genes than those in DS+NH controls. Gene set enrichment tests suggested an enrichment of large deletions intersecting ciliome genes. Most importantly, the scale of this study showed that, even in the sensitized DS population, there are no large, common CNVs with a major effect on AVSD that could account for the 2,000-fold increased risk in DS. We have also shown that common SNPs cannot account for the increased risk of CHD in this same cohort (Ramachandran et al. 2015).

In the current study, we focus specifically on CNVs on chromosome 21 and test two primary hypotheses: 1) individuals with DS carrying chromosome 21 deletions may be protected from AVSD, because these deletions return AVSD susceptibility loci back to disomy, and 2) individuals with DS carrying chromosome 21 duplications are at increased risk for AVSD, because these duplications boost the dosage of AVSD susceptibility loci beyond a tolerable threshold. In addition to testing these hypotheses, we used our large cohort of 198 DS+AVSD cases and 211 DS+NH controls in an independent replication of Sailani et al. (2013). They screened for CNVs on chromosome 21 in a similarly defined DS cohort of 55 DS+AVSD cases and 53 DS+NH controls and reported two common CNVs significantly associated with AVSD.

## Results

We used rigorous quality control (see METHODS) to identify deletions and duplications on the trisomic chromosomes 21 in 409 DS individuals, including 355 Caucasians (174 DS+AVSD cases and 181 DS+NH controls) and 54 African Americans (24 DS+AVSD cases and DS+NH 30 controls). This analysis revealed a high-quality set of 215 individual deletions and 59 individual duplications (Table 1). For Caucasians and African Americans, respectively, 91% and 100% of these deletions had 50% reciprocal overlap with deletions in the Database of Genomic Variants (DGV) (MacDonald et al. 2014, http://dgv.tcag.ca). For duplications, 82% and 60% of these variants were reported in the DGV.

**Table 1.** CNV summary statistics for cases and controls stratified by race/ethnicity. Cases (DS+AVSD) are defined as those with Down syndrome and complete atrioventricular septal defect. Controls (DS+NH) are individuals with Down syndrome without a congenital heart defect.

|  |  | # Participants<br>(male/female) | Type | # Variants | Average<br>per person | Size range<br>(kb) | Median size<br>(kb) |
| --- | --- | --- | --- | --- | --- | --- | --- |
| **Caucasians** | **Cases**<br>**DS+AVSD** | 77 / 97 | Deletions | 66 | 0.38 | 1.8 - 159.2 | 10.7 |
|  |  |  | Duplications | 31 | 0.18 | 11.6 - 395.1 | 23.1 |
|  | **Controls**<br>**DS+NH** | 104 / 77 | Deletions | 71 | 0.39 | 1.8 - 239.1 | 10.7 |
|  |  |  | Duplications | 23 | 0.13 | 10.4 - 200.0 | 18.0 |
| **African**<br>**Americans** | **Cases**<br>**DS+AVSD** | 6 / 18 | Deletions | 36 | 1.50 | 2.1 - 44.1 | 4.4 |
|  |  |  | Duplications | 1 | 0.04 | 491.9 | 491.9 |
|  | **Controls**<br>**DS+NH** | 18 / 12 | Deletions | 42 | 1.40 | 2.1 - 260.3 | 7.2 |
|  |  |  | Duplications | 4 | 0.13 | 4.5 - 14.4 | 10.9 |

## No single CNV of large effect is associated with AVSD in DS

We performed association testing of single deletion and duplication regions along chromosome 21, as well as of single genes intersected by deletions or duplications, controlling for possible population stratification (see Methods). Though in Caucasians we had 80% power to detect risk variants of 5% allele frequency with an odds ratio of 2.2 or greater (alpha level of 0.05), no single CNV region was associated with AVSD (Supplemental Fig. 1). We also tested for association of single genes with any intersection by CNVs and found no suggestive association. With our small African American cohort, we had 80% power to detect a risk CNV with an odds ratio of 6.3 at an allele frequency of 0.05 and an alpha level of 0.05. Again, we found neither a single CNV nor any CNV-intersected gene on chromosome 21 associated with AVSD in our Down syndrome population.

**Figure 1.**
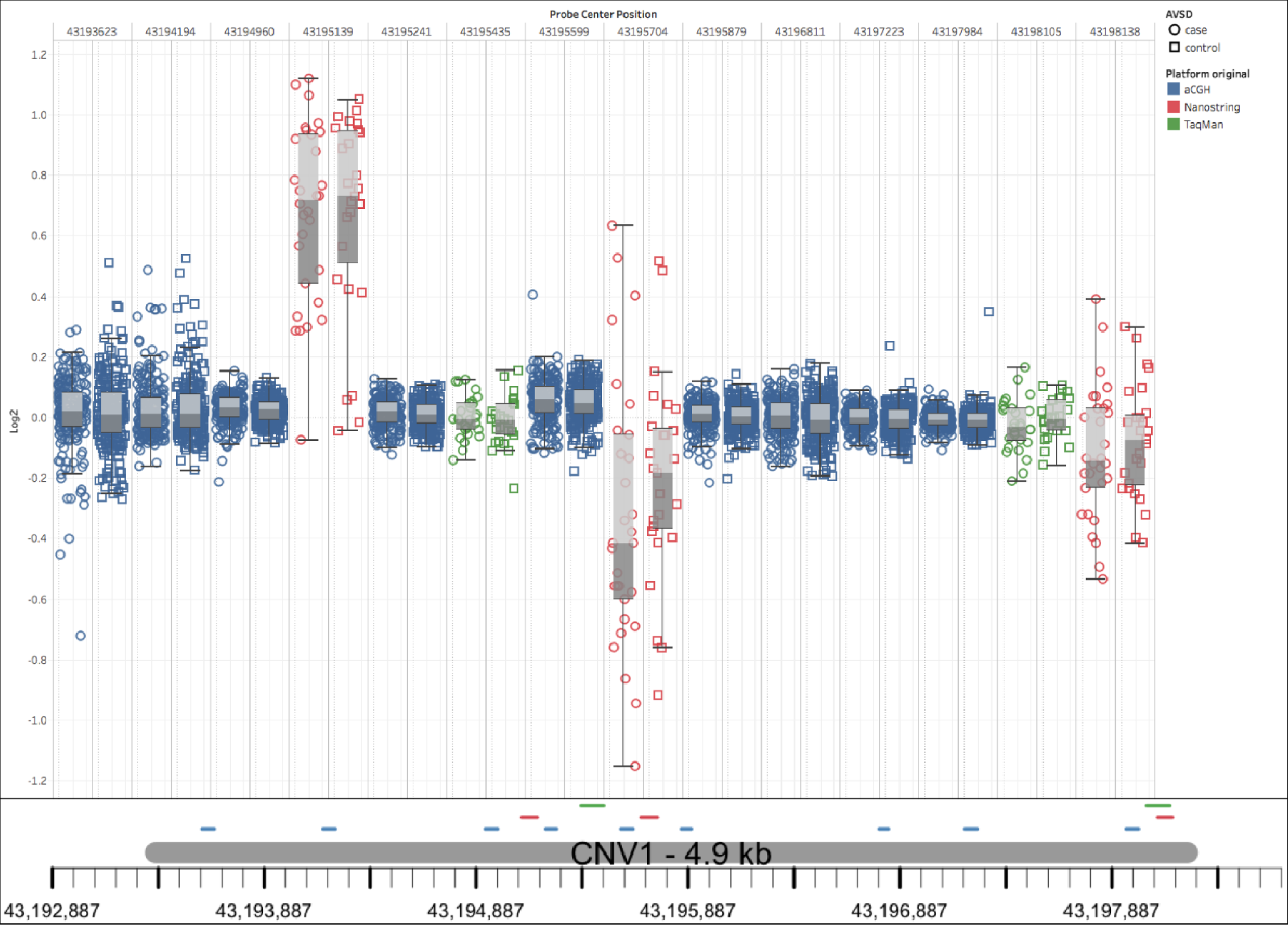
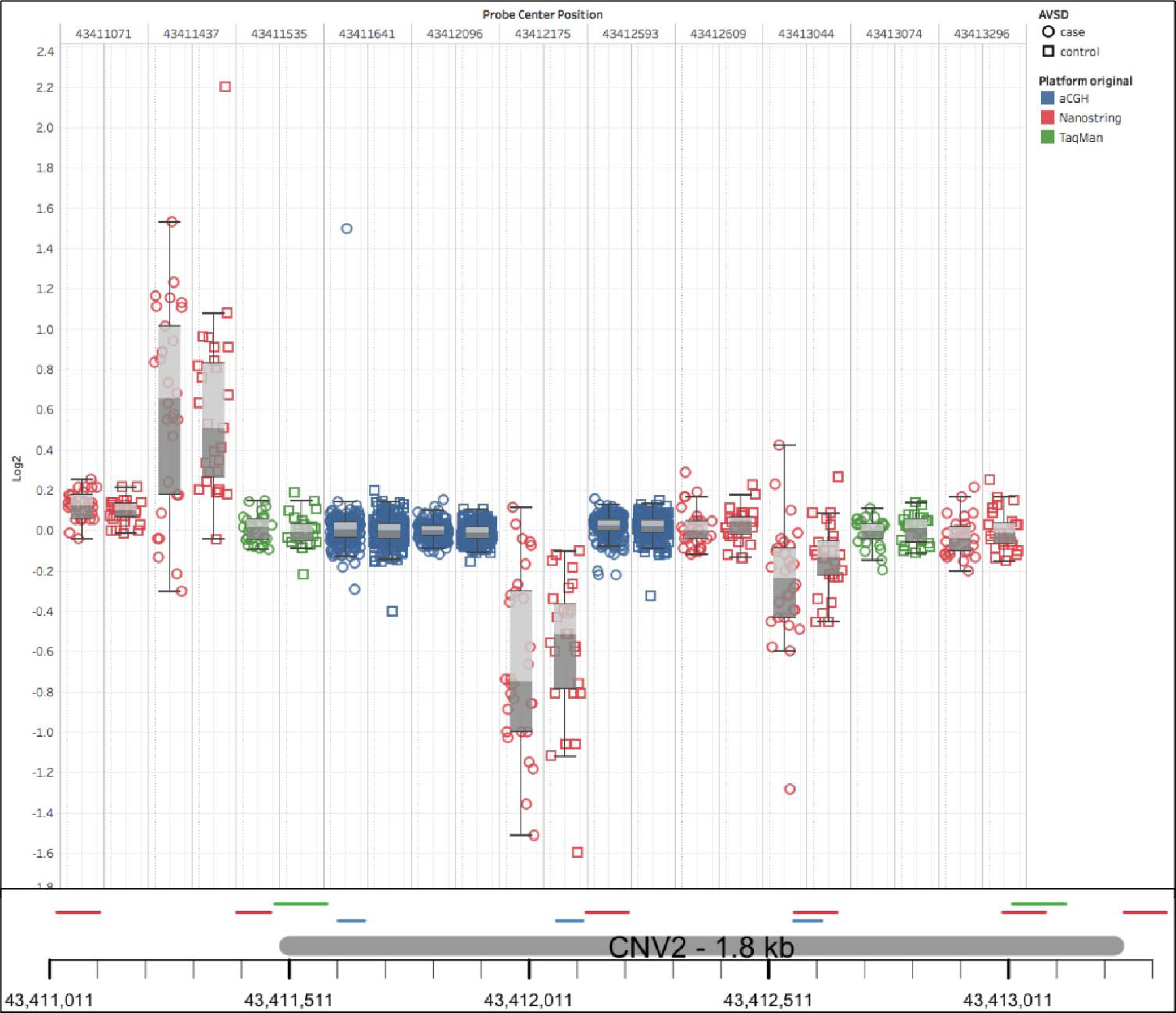
Summary results from analyses to attempt to replicate previously reported DS+AVSD-associated common CNVs with three technologies. Boxplots show inner quartile range of log2 ratios for tested DS samples with whiskers reaching the minimum and maximum observed ratios. Case values are in filled boxes, and control values are in outlined boxes. CGH probes are blue, NanoString probes are red and TaqMan® probes are black. Boxplots are ordered by genomic location, and their precise locations are indicated beneath the plots. Neither CGH probes nor TaqMan® Copy Number assays detected aberrant copy numbers or differences between cases and controls. Varying results were found across these loci by NanoString probes, with some probes showing differences in log2 means between cases and controls. Within these same proposed small CNV loci, NanoString probes called all possible combinations of copy gain, loss, and no change within the same small cohort. When compared to adjacent CGH and TaqMan® probes, it is clear the NanoString probes are not reliable predictors of copy number state at this locus.

## Burden of chromosome 21 deletions

We tested our first hypothesis that individuals with DS carrying chromosome 21 deletions may be protected from AVSD, because these deletions return AVSD susceptibility loci back to disomy. To do this, we compared the “burden” of chromosome 21 deletions among DS+AVSD cases to that of DS+NH controls. Using PLINK v1.07 (Purcell et al. 2007), we determined whether there was a greater average number of deletions per person measured in two ways: 1) as an increased number of bases covered by deletions and 2) as an increased average number of genes intersected by deletions on chromosome 21. We tested all deletions, filtered by allele frequency (common ≥0.01 or rare <0.01), and whether they were reported in the DGV. Our analyses revealed no effect of deletions providing a protective effect against AVSDs in Caucasians (Supplemental Table 1). In contrast, African American DS+NH controls were significantly more likely to have more bases covered by deletions within the full deletion set (average total bases covered by deletions: 33.45 kb in DS+NH controls vs. 13.06 kb in DS+AVSD cases; empirical p-value = 0.04; Supplemental Table 1). When we filtered deletions by frequency, we found that this effect in African Americans was driven by rare variants of less than 1% frequency in our study sample: African American DS+NH controls with rare deletions have on average 45.63 kb covered by rare deletions versus 12.8 kb in DS+AVSD cases (empirical p-value = 0.02, Supplemental Table 1>).

To further test our hypothesis, we redefined our definition of CNVs that might reduce a gene to disomy by disrupting gene function by including: 1) deletions that intersected an exon and 2) duplications that intersected an exon, but did not envelope an entire gene. In African Americans, this reduced the set to only two CNVs, a deletion and duplication in two controls (1-sided Fisher’s exact p-value = 0.32). In Caucasians, this produced a set of 41 CNVs in DS+AVSD cases and 41 CNVs in DS+NH controls that reduce a gene back to disomy; thus, there was no indication that individuals with DS without heart defects are protected by CNVs that reduce a gene back to disomy (Supplemental Table 3).

## Burden of chromosome 21 duplications

To test our second hypothesis that chromosome 21 duplications increase the risk for AVSD, we compared the burden of chromosome 21 duplications among DS+AVSD cases compared with DS+NH controls. In Caucasians, a number of findings were consistent with this hypothesis (Supplemental Table 2). We observed that duplications, on average, affect more bases in cases (83.53 kb) than in controls (40.49 kb) (empirical p-value = 0.09). Caucasian cases also had twice the rate of genes duplicated compared with controls (0.22 in cases versus 0.10 in controls; empirical p-value = 0.07). Rare CNVs in Caucasians drive these effects. For example, cases had a higher rate of rare duplications than controls (0.09 in cases versus 0.03 in controls; empirical p-value = 0.06). More specifically, cases have five times the rate of genes intersected by rare duplications (0.16) compared with controls (0.03) (empirical p-value = 0.04). These effects remain by filtering for variants not in the DGV, as they are all rare variants. Given the low number of duplications in the African American samples, we did not see an increased burden of duplications among DS+AVSD cases (Supplemental Table 2).

**Table 2.** Counts of cilia genes on chromosome 21 intersected by CNVs. Gene Set Enrichment Analysis (GSEA) tests for enrichment of CNVs in 19 cilia genes compared to all other genic CNVs. Two cilia genes are intersected by deletions in controls, which suggests an enrichment from a GSEA permuted p-value of 0.1. We find the opposite pattern in duplications where two genes are intersected only in controls (GSEA permuted p-value of 0.2). Although not statistically significant, the pattern is consistent with our stated hypothesis.

|  | Number of cilia genes intersected |  | p-value |
| --- | --- | --- | --- |
|  | Cases<br>DS+AVSD | Controls<br>DS+NH |  |
| Deletions | 0 | 2 | 0.1 |
| Duplications | 2 | 0 | 0.2 |

Again, to further test this hypothesis, we filtered duplications for those that contained a full gene and found six in cases and one in a control (one-sided Fisher’s exact p-value = 0.10; odds ratio = 5.3, and 95% C.I. = 0.75-Infinity). Two of these duplications reside in the same case individual.

## Gene Set Enrichment and Gene Ontology Analyses

Previous reports suggest that genetic variation in cilia genes plays a role in AVSD in DS (Ripoll et al. 2012; Ramachandran et al. 2015; Burnicka-Turek et al. 2016). There are 19 genes on chromosome 21 implicated as part of the ciliome (Supplemental Table 9, McClintock et al., 2008). We performed Gene Set Enrichment Analysis (GSEA) with PLINK v1.07 (Purcell et al. 2007) to test two hypotheses: 1) chromosome 21 deletions are more likely to intersect cilia genes than other genes in DS+NH controls and 2) chromosome 21 duplications are more likely to intersect cilia genes than other genes in DS+AVSD cases. In Caucasians, two putative cilia genes, *DYRK1A* and *PDXK,* were intersected by deletions in controls; no cilia genes were intersected by deletions in cases (Table 2). While the counts are low, this finding is suggestive (permuted p-value = 0.1) by GSEA. Similarly, for duplications, we observed the inverse: duplications intersect with two cilia genes, *USP25* and *ITSN1*, in cases and none in controls (permuted p-value = 0.2). When we combined deletions and duplications that disrupt exons and thus reduce that gene to disomy, we found one case and two controls with a CNV disrupting a cilia gene exon (permuted p-value = 0.25).

To uncover novel pathways disrupted by CNVs in DS-associated AVSD, we performed a Gene Ontology analysis with the ClueGO v2.2.5 (Bindea et al. 2009) plugin in Cytoscape v3.3.0 (Shannon et al. 2003), providing lists of genes that were intersected by deletions and duplications only in cases or only in controls (Supplemental Table 4). As we assume the same pathways leading to AVSD will be perturbed in all humans, we combined lists of genes from African Americans and Caucasians, creating eight gene lists: 1. genes intersected by deletions only in cases or 2. only in controls; 3. genes intersected by duplications only in cases or 4. only in controls; 5. genes with an exon intersected by a deletion or non-gene-enveloping duplication only in cases or 6. only in controls; and 7. genes completely duplicated only in cases or 8. only in controls. In the deletion gene lists, only those within DS+NH controls clustered into a pathway. Deleted genes in controls were overrepresented in protein heterotrimerization (GO:0070208, p-value = 0.0002). In the duplication gene lists, significant pathway enrichment was found only in DS+AVSD cases. These duplication-intersected genes were significantly enriched for synaptic vesicle endocytosis (GO:0048488, p-value = 0.0001). While genes with exons disrupted by CNVs in cases showed no enrichment in biological pathways, those in controls were enriched in protein heterotrimerization (GO:0070208, p-value = 0.0047). There was only one gene and one ncRNA completely duplicated in controls, while there were 19 genes duplicated in cases, and they were enriched in the process of histone methylation (GO:0016571, p-value = 0.0017).

## Replication of previous findings of common CNVs associated with DS-associated AVSD

We next sought to replicate two loci previously reported to be significantly associated with AVSD in a collection of individuals with DS (Sailani et al. 2013). CNV1 at chr21:43,193,374-43,198,244 (hg19) was observed as a deletion in 18% of the 55 cases with DS+AVSD and 0% of the 53 DS+NH controls and as a duplication in 7% of cases and 0% in controls. CNV2 at chr21:43,411,411-43,413,231 was found as a deletion in 24% of controls versus 0% in cases and as a duplication in 14% of cases versus 11% of controls (Table 3). In an internal validation study based on 49 DS+AVSD cases and 45 DS+NH controls, Sailani et al. used NanoString nCounter® technology and found significant differences in copy number ratios in probes targeting these loci.

**Table 3.**
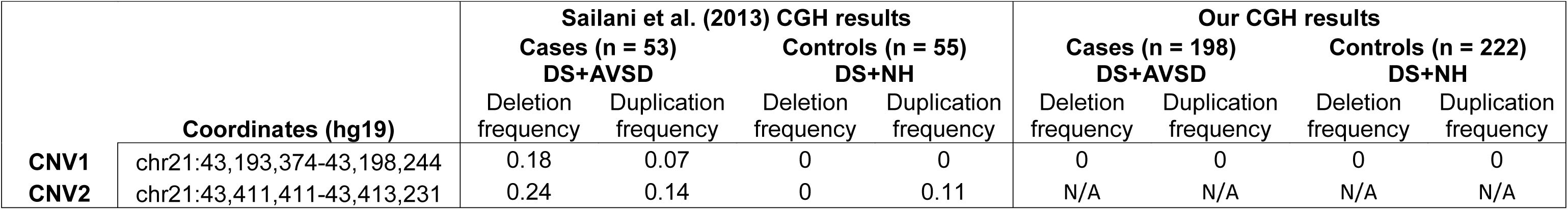
Comparison of DS+AVSD significantly associated CNVs from Sailani et al. (2013) to our current study. We did not replicate the previously reported significant association of common deletions and duplications at CNV1. Our CGH array did not have at least six probes inside CNV2 and thus was undetectable by our methodology.

Our aCGH experiments had 19 probes within CNV1 and did not detect any CNV, though our sample size is four times larger than that of Sailani et al. (Table 3). Our custom array had only three probes in the CNV2 locus, and thus we were not able to detect it with our stringent criteria, which required six or more probes to call a CNV. We performed a NanoString experiment on a subset of our sample (49 cases and 45 controls) using the same CodeSet probes that Sailani et al. found to be significantly associated with DS+AVSD. We followed the same methodology they used to analyze the NanoString data. For each probeset, a ratio of the probe’s copy number count from a test individual over that of a reference DS sample was computed. A Mann-Whitney U-test was then applied at each probe, testing for a difference in the mean copy number count ratios between cases and controls. In CNV1, one of the three probes found significant in the Sailani et al. sample, we found a significant difference in nCounter copy number ratios between our cases and controls (p-value = 0.007) (Table 4). At CNV2, two of the six probes found significant in Sailani et al. were marginally significant in our dataset (p-values = 0.054 and 0.056).

**Table 4.**
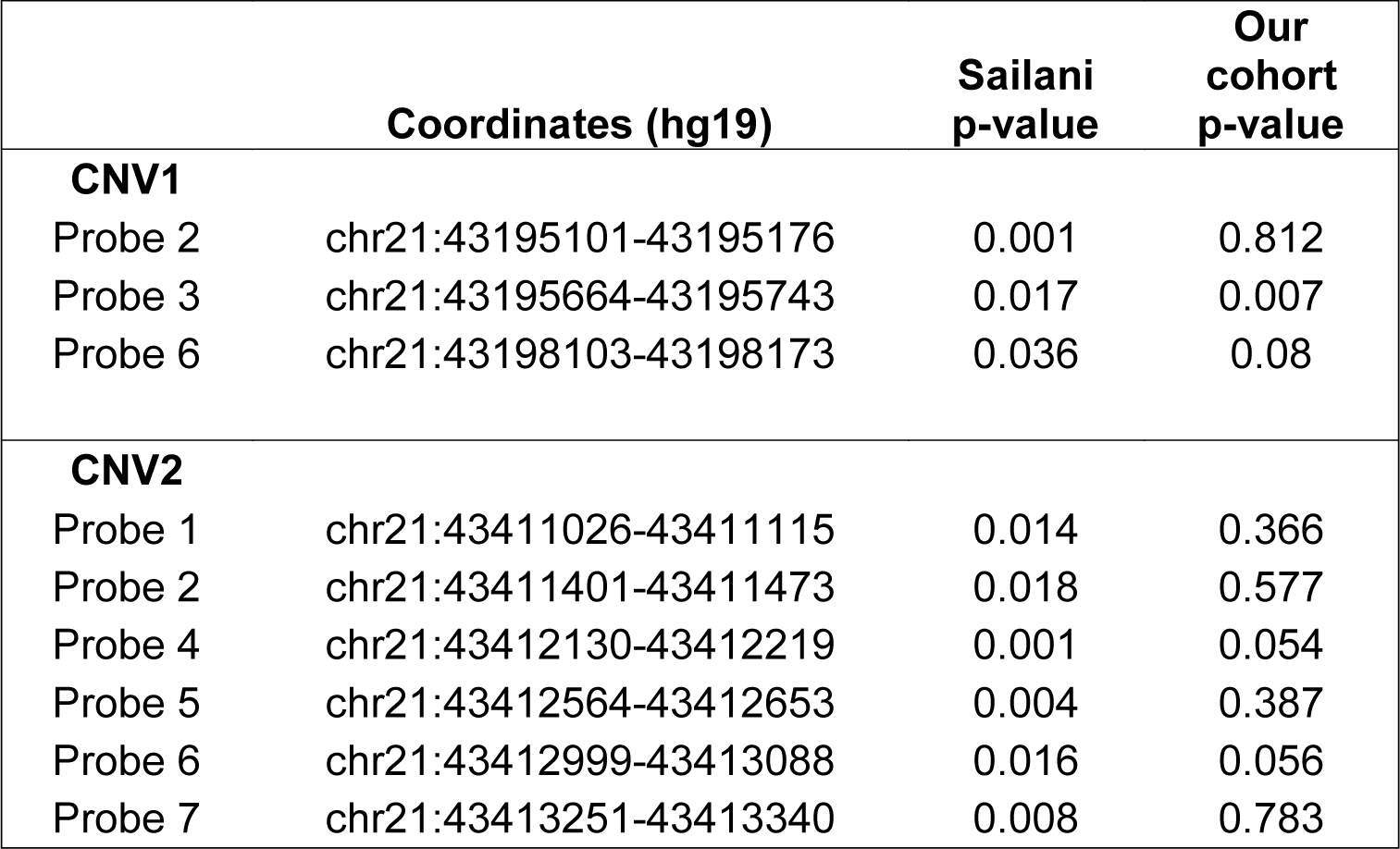
Comparison of DS+AVSD significantly associated CNVs from Sailani et al. (2013) to our current study using NanoString technology. We performed NanoString nCounter assays on 46 DS+AVSD cases and 45 DS+NH controls using the same probes used by Sailani et al. (2013) in their CNV replication experiment that included 49 cases and 45 controls. To maintain congruency, we applied their assessment strategy to test for mean differences in normalized count (CN) ratios using a one-sided Mann-Whitney U-test. In CNV1, we detected a significant difference in CN ratios for probe 3 (using Sailani et al. nomenclature), but did not find this relationship for the other two probes previously found significant by Sailani et al. (2013). In CNV2, two of the six previously significant probes were marginally significant in our experiment.

|  | Coordinates (hg19) | Sailani<br>p-value | Our<br>cohort<br>p-value |
| --- | --- | --- | --- |
| **CNV1** |  |  |  |
| Probe 2 | chr21:43195101-43195176 | 0.001 | 0.812 |
| Probe 3 | chr21:43195664-43195743 | 0.017 | 0.007 |
| Probe 6 | chr21:43198103-43198173 | 0.036 | 0.08 |
| **CNV2** |  |  |  |
| Probe 1 | chr21:43411026-43411115 | 0.014 | 0.366 |
| Probe 2 | chr21:43411401-43411473 | 0.018 | 0.577 |
| Probe 4 | chr21:43412130-43412219 | 0.001 | 0.054 |
| Probe 5 | chr21:43412564-43412653 | 0.004 | 0.387 |
| Probe 6 | chr21:43412999-43413088 | 0.016 | 0.056 |
| Probe 7 | chr21:43413251-43413340 | 0.008 | 0.783 |

The mixed results within NanoString and aCGH experiments led us to assess the validity of these findings with a third technology (Fig. 1). Two TaqMan® probesets were selected within each CNV and tested in 46 DS+AVSD cases and 46 DS+NH controls, including the same cases and controls analyzed with NanoString. No deletions were identified by either probeset in CNV1 or CNV2 (Table 4). At CNV1, a duplication was detected in one individual (DS+NH control) by one probeset; the other did not detect a copy number change. At CNV2, a duplication was detected in one individual (a DS+NH control) by both probesets. No DS+AVSD cases had copy number calls at either CNV.

## Discussion

Our cohort of individuals with DS with complete AVSD and those with structurally normal hearts represents the largest study of its kind to date. In addition, our CNV dataset was built applying conservative quality control metrics on probe, array, and sample inclusion, yielding robust conclusions after analysis. The composite set of CNVs, which required concordance between two well-established CNV calling algorithms, generated a dataset with a low likelihood of false-positive findings, as indicated by their high representation in the DGV (94% of deletions and 80% of duplications).

Given our large sample size for this relatively rare condition, we had 80% power to detect a single CNV at 5% population frequency with an odds ratio of 2.2 or greater in Caucasians and 6.3 in African Americans (empirical p-value of 0.05). We detected no single CNV with an effect size of this magnitude. Our data suggest it is unlikely for a single common variant larger than 1.7 kb on chromosome 21 to explain the 2,000-fold increased risk for AVSD on a trisomy 21 background. This is consistent with our previous findings (Ramachandran et al. 2015; Ramachandran et al. 2015).

We specifically examined genes required for proper cilia function because of the mounting evidence for their involvement in CHD. We found a suggestive association of perturbations in such genes, consistent with our previous genome-wide results in this same cohort, where we found an increased burden of deletions overlapping cilia genes in cases versus controls (Ramachandran et al. 2015). Other support for ciliome involvement comes from multiple lines of evidence. In a forward genetic mouse screen, 87,355 fetuses from mutagenized mice resulted in 218 mice with CHDs (Li et al. 2015). Exome sequencing of 113 of these mice revealed 91 recessive mutations in 61 genes, of which 34 were involved in cilia function. Disruption of cilia-related genes is also linked to AVSD in lymphoblastoid cell lines (LCLs) from individuals with DS (Ripoll et al. 2012). Comparing gene expression profiles of LCLs from individuals with DS without CHDs to those with atrial septal defects (ASDs), ventricular septal defects (VSDs), or AVSDs, they found significant deregulation of cilia genes within the AVSD group. Furthermore, principal component analysis of the expression profiles separate individuals with AVSDs from those with ASDs or VSDs, pointing to a different etiology of disease progression and substantiating the need to study phenotypically distinct CHDs independently. Most recently, Burnicka-Turek et al. characterized independent mouse lines with deleterious non-synonymous mutations induced in the cilia genes *Dnah11* and *Mks1* that caused AVSD (Burnicka-Turek et al. 2016).

We were unable to replicate two previously reported common CNVs on chromosome 21 associated with DS+AVSD in a smaller cohort by Sailani et al. (2013). Although our custom array did not have enough probes to reliably detect CNV2, we were powered to detect CNV1 and did not find this CNV in either cases or controls in our larger population. Sailani et al. replicated their finding in an independent sample by reporting differences in means of NanoString nCounter® probe ratios between cases and controls. At CNV1, we tested three of these significant probesets with available coordinates and found only one to be significant by Mann-Whitney U-test. At CNV2, we tested the six probes previously found significant and found two of the six to be marginally significant. Sailani et al. calculated ratios of probe counts of the test sample over that of the reference sample and tested for ratio differences between cases and controls using a one-sided Mann-Whitney U-test. We applied their techniques to a similar sized subset of our larger cohort and found inconclusive results to support their findings. As a final validation of these proposed DS+AVSD-associated CNVs, we performed TaqMan® Copy Number assays with two probe sets for each CNV. As detailed in Results, no deletions were detected at CNV1 or CNV2. Thus, we failed to replicate their reported findings for CNV1 in a cohort that was four times as large and for both CNV1 and CNV2 using a similar technology and a follow-up gold-standard technology (Fig. 1). We conclude that the associations to DS+AVSD of CNV1 and CNV2 are likely false positives.

Our study stands in agreement with the consensus of other studies reporting complex heterogeneity of genetic contributions to atrioventricular septum and valve development in both the disomic population and in individuals with trisomy 21 (Robinson et al. 2003; Ackerman et al. 2012; Al Turki et al. 2014; Ramachandran et al. 2015; Ramachandran et al. 2015; Priest et al. 2016). Our data support the nuanced hypotheses that deletions on chromosome 21 on a trisomic background reduce the risk for AVSD and duplications on chromosome 21 further increase risk of AVSD in DS. These effects were enriched when considering rare (MAF<0.01) deletions and duplications. Rare deletions have been previously implicated within our DS cohort, where DS+AVSD cases were found to have a greater genome-wide burden of rare, large (>100 kb) deletions (Ramachandran et al. 2015; Ramachandran et al. 2015)

Moving forward, genetic studies of CHD in DS, as well as nonsyndromic CHDs, should be designed with this considerable genetic heterogeneity in mind. It is clear that, while trisomy 21 alone increases the risk for AVSD 2,000 fold, its probable mode of action is through epistatic interactions among many genes, at least some of which are necessary for the structure and function of cilia. Untangling these complex risk factors will require a larger cohort of individuals with DS with and without CHDs to find susceptibility loci of measurable effect. As these cohorts continue to grow, efforts should focus on exome and whole-genome sequencing approaches that identify rare variants, whose effects can be tested for burdening candidate genetic pathways of cardiogenesis. Finally, environmental factors require greater consideration, and resources should be prioritized to gather broad epidemiological data and link them to genomic resources.

## Methods

### DNA samples

Participant samples were collected as described previously (Freeman et al. 1998; Freeman et al. 2008; Locke et al. 2010; Ramachandran et al. 2015; Ramachandran et al. 2015). Individuals diagnosed with full or translocation trisomy 21, documented by karyotype, were recruited from centers across the United States. Institutional review boards at each enrolling institution approved protocols, and informed consent was obtained from a custodial parent for each participant. A single cardiologist (K. Dooley) identified cases from medical records as individuals with a complete, balanced AVSD diagnosed by echocardiogram or surgical reports (DS+AVSD). Controls were classified as individuals with a structurally normal heart, patent foramen ovale, or patent ductus arteriosus (DS+NH).

Genomic DNA was extracted from LCLs with the Puregene DNA purification kit according to the manufacturer’s protocol (Qiagen, Valencia, CA). DNA quantity and quality were checked on a NanoDrop ND-1000 spectrophotometer (NanoDrop Technologies, Wilmington, DE) and assessed for integrity on 0.8% agarose gels stained with ethidium bromide.

### Microarray design and processing

All analyses used the human genome reference hg19 build. We designed a custom 8x60k Agilent (Agilent Technologies, Santa Clara, CA) CGH array using eArray (https://earray.chem.agilent.com accessed April, 2011). The array consisted of 52,944 60-mer DNA probes targeting human chromosome 21, providing a mean spacing of 673 bp and a median spacing of 448 bp, as well as a genomic backbone of probes and Agilent’s control probes (design file: ADM2Chr21_60k_final_033839_D_F_20120731.xml).

Array hybridization was processed according to Agilent’s protocol and scanned on an Agilent SureScan High-Resolution Microarray Scanner at Emory University. A single female (GEO accession individual ID: 246) with trisomy 21 and no CHD was used as the reference sample for all test individuals. This individual had a known deletion at chr21:45,555,257-45,615,042, which would be detected as a duplication in all test samples.

### Sample quality control

We performed aCGH on a total of 550 DS samples. We preformed three stages of sample/array quality control (QC). We first performed Agilent’s recommended QC. Their recommended QC cutoff for arrays is a derivative log_2_ ratio (DLR) <0.3. DLR is a measure of probe-to-probe noise and is the standard deviation of adjacent probe log_2_ differences. Twenty-five samples failed to meet this threshold and were excluded (Supplemental Fig. 2).

Second, while the remaining 525 microarrays met Agilent’s basic QC parameter of DLR <0.3, visual inspection of log_2_ plots revealed a number of arrays with an increased probe variance. To quantitatively assess and account for this effect, we calculated the variances of intra-array probe log_2_ ratios to develop a conservative array inclusion criterion. We excluded 74 arrays with variance ≥1 standard deviation (SD) over the mean from any further analysis (Supplemental Fig. 3).

Third, to avoid biasing an individual microarray toward over- or undercalling gains or losses, it is important that the mean log_2_ ratio across the array is near the expected value of zero. The means of the intra-array probe log_2_ were calculated on the 451 remaining arrays (grand mean = -0.00045), and 25 arrays with individual means ≥2 SD from the group mean were removed (Supplement Fig. 4). After CNV detection (described below), we removed clear outlier samples that had the number of CNVs (deletions or duplications) called >5 SD over the mean. This eliminated five samples.

To avoid spurious association results based on population stratification, we performed principal component analysis on most of our samples that had genome-wide SNP data available from our previously published study (Ramachandran et al. 2015). Four samples without genotyping data were removed from further CNV analyses. In PLINK (version 1.9; Chang et al. 2015), SNPs were removed that had >10% missingness or failed the Hardy-Weinberg equilibrium exact test with a p-value <1x10^−6^. Common SNPs with minor allele frequency >0.05 were pruned by PLINK’s “--indep-pairwise” command within 50-kb windows, a five SNP step, and an r^2^ threshold of 0.2, leaving 552,943 SNPs. The first five eigenvectors were calculated using the R package SNPRelate (Zheng et al. 2012) and plotted (Supplemental Figs. 5-9). PCA round 1 clearly separated self-identified African Americans from Caucasians. Round 2 was performed separately on African Americans and Caucasians. Six African Americans were visibly clear outliers (Round 2 PC1 ≤-0.127) and were removed from further analyses (Supplemental Figs. 6 and 8). Five Caucasians were clear outliers (Round 2 PC1 ≤-0.1) and were also removed (Supplemental Figs. 7 and 9). The final cohort contained 198 cases and 211 controls (Supplemental Table 6).

### CNV calling

We also evaluated the quality of data at the probe level. Because custom CGH arrays contain probes with unpredictable binding characteristics, the variances of normalized probe fluorescent signals were calculated, and 2,193 probes with inter-array variance ≥1 SD above the mean were removed (Supplemental Fig. 10). These calculations were done on the full set of arrays passing the above DLR criteria and before the above intra- and inter-array probe log2 variance calculations and filtering.

We used two algorithms, ADM2 and GADA, to identify putative CNVs (Pique-Regi, R et al. 2010). We required that CNVs be called by both algorithms to be included in the analysis. Parameters for Agilent’s ADM2 algorithm were set within their Genomic Workbench software (version 7.0.4.0) as follows: ≥6 probes, average log_2_ shift of ±0.2, use of the diploid peak centralization, 2-kb window GC correction, intra-array replicates combined, and Fuzzy Zero applied. GADA adjustable parameters are the minimum probe number for a CNV to be called (MinSegLen) and a threshold, T_m_, referring to the minimum t-statistic that a predicted breakpoint must reach during its Backward elimination procedure. We empirically optimized the GADA T_m_ variable across a range of 4.5 to 20.5, by half steps, and evaluated performance based on two criteria: 1. Whether the algorithm detected duplications in at least 80% of our test samples at our known reference deletion, and 2. Whether the algorithm detected common deletions found in the 1000 Genomes’ Phase 3 release of structural variants at a similar population frequency.

Compressed .vcf data and accompanying .tbi file for chromosome 21 produced from whole-genome sequencing by the 1000 Genomes Consortium was downloaded from http://hgdownload.cse.ucsc.edu/gbdb/hg19/1000Genomes/phase3/ on Feb. 27^th^, 2016. Tabix commands created unzipped .vcf files covering chr21:13000000-47000000, and variants denoted ‘SVTYPE’ were filtered with shell commands, and variants denoted as deletion, duplication, or both (multi-allelic) were analyzed. The lower limit of CGH detection was set at ≥1798 bp, and variants of less than 1,798 bp were removed from the 1000 Genomes’ comparison set.

Of seven common CNVs on chromosome 21, our CGH array had at least six probes in two of these variants: esv3646598 and esv3646663. esv3646598 has a frequency of 0.064 in individuals of European ancestry (0.004 in African ancestry). esv3646663 has a frequency of 0.227 in individuals of African ancestry (0.001 in Europeans). We do not call absolute copy number from CGH data, and a deletion can represent zero, one, or two copies of the three expected. Thus, an upper frequency bound was set as Freq = (3(d)+0(N-d)) / 3(N), while the lower bound was set as Freq = (1(d)+0(N-d)) / 3(N), where d equals the number of times the deletion was called and N equals the total number of chromosomes. For both variants, in their respective ancestral population, T_m_ = 8 detects common structural variants within the expected range and also maximizes the detection of our reference deletion (Supplemental Figs. 11 and 12). GADA was then launched using a custom R script applying the following parameters: estim.sigma2 = TRUE, MinSegLen = 6 and T_m_ = 0.8.

CNVs ≥1 Mb were removed (14 deletions; 7 duplications) after visually checking log_2_ plots to confirm these were likely false positives. Variants with breakpoints inside our reference deletion (chr21:45555257-45615042) were removed (0 deletions; 354 duplications). The p-arm and pericentromeric region of chromosome 21 are poorly mapped, and variants with breakpoints inside chr21:0-15400000 were removed (2 deletions; 1 duplication). Clear outliers containing large numbers of deletions or duplications were removed. We used a threshold of >5 SD over the mean of 0.73 deletions and 0.15 duplications calculated among the 426 arrays. Five SDs over the mean corresponded to more than five deletions or two duplications within one array. These five samples contained 81 deletions and six duplications. The final dataset includes 215 deletions and 59 duplications, of which 92% of deletions and 73% of duplications have 50% reciprocal overlap with variants in the (DGV), indicating a low rate of false positives (Supplemental Table 7).

### Replication study of Sailani et al. (2013)

Two common CNVs were found to be associated with DS+AVSD in the study by Sailani et al. (2013). To try to replicate that finding, we used identical NanoString probes in 96 cases and controls from our DS cohort. We included probes that showed significant copy number differences between their cases and controls, totaling four of the eight probes for CNV1 and five of the seven probes for CNV2 (Supplemental Table 8). Samples were processed by the Gene Expression Analysis Laboratory at The University of Tennessee. Additionally, two TaqMan® (Applied Biosystems, Grand Island, NY) assays targeting each locus were selected for CNV1 and CNV2 (Supplemental Table 9). These assays were performed by the Emory Integrated Genomics Core on the same 96 samples tested by NanoString. Copy number calls were made by TaqMan’s CopyCaller^TM^ software, and calls with a confidence probability less than 0.8 were dropped.

### CNV association and burden analyses

We used PLINK v.1.07 to carry out association and burden analyses separately for deletions and duplications. To explicitly test the hypothesis that a gene reduced to two functional copies provides protection against AVSD in DS, we combined deletions that intersect an exon (refGene-hg19 updated July 3, 2016) with duplications that have predicted breakpoints within an exon to form a “reduced to disomy” set of CNVs. We also explicitly tested the inverse hypothesis: that genes entirely duplicated increase the risk for AVSD in DS. Three testing paradigms were performed: 1) burden analyses using the --cnv-indiv-perm and --cnv-count commands, 2) associations with individual CNV regions using --cnv-count, and 3) associations with individual genes overlapped by a CNV --cnv-intersect and --cnv-test-region. Empirical p-values of significance were determined by performing one million permutations for each test. These p-values are one-sided, and we tested for excess burden of duplications in cases and for deletions in controls. These three testing paradigms were applied to the full dataset, as well as subsets of CNVs filtered by overlap in the Database of Genomic Variants (downloaded January, 2016), and by CNV frequency of greater than or less than 1% within out study population. Burden analysis in PLINK tests for differences between cases and controls using three different approaches: 1) Is there a difference between the average number of CNVs per person (RATE)?; 2) Is there a difference in the average number of bases covered by all CNVs (KBTOT)?; and 3) Is there a difference in the average number of genes intersected by CNVs per person (GRATE)? We performed burden tests across deletions and duplications on chromosome 21 as entire sets and filtered by the allele frequency of the CNV (common or rare <0.01) and by their existence or lack thereof in the Database of Genomic Variants.

### Gene Set Enrichment and Gene Ontology Term Analyses

We used PLINK v1.07 to perform Gene Set Enrichment Analysis (GSEA) (Raychaudhuri et al. 2010) on a set of cilia-related genes (Table 15) compiled by McClintock et al. 2008 and previously implicated in Down syndrome-associated AVSD (Ripoll et al. 2012; Ramachandran et al. 2014; Li et al. 2015). GSEA tests the hypothesis that cilia genes are enriched for CNVs compared to all chromosome 21 genic CNVs and assigns a one-sided empirical p-value by one million permutations, indicating positive enrichment in cases for duplications or in controls for deletions.

We performed a Gene Ontology (GO) term analysis using the ClueGO Extension in Cytoscape v.3.3.0 (Bindea et al. 2009). Gene lists were created based on those intersected by deletions or duplications only in cases or only in controls (Supplemental Table 3). Each gene set was analyzed with the following settings: Ontologies = Go-Biological Process downloaded May 5^th^, 2016, Evidence = All, Pathway Significance = 0.05, GO Tree Interval = 3 minimum level and 13 maximum level, two gene minimum or 1% of pathway genes, and Bonferroni step-down p-value correction with mid-P-values. Other parameters were left at default settings.

